# Controlling Cell-Free Gene Expression Behavior by Tuning Membrane Transport Properties

**DOI:** 10.1101/604454

**Authors:** Patrick M. Caveney, Rosemary M. Dabbs, William T. McClintic, S. Elizabeth Norred, C. Patrick Collier, Michael L. Simpson

## Abstract

Controlled transport of molecules across boundaries for energy exchange, sensing, and communication is an essential step toward cell-like synthetic systems. This communication between the gene expression compartment and the external environment requires reaction chambers that are permeable to molecular species that influence expression. In lipid vesicle reaction chambers, species that support expression – from small ions to amino acids – may diffuse across membranes and amplify protein production. However, vesicle-to-vesicle variation in membrane permeability may lead to low total expression and high variability in this expression. We demonstrate a simple optical treatment method that greatly reduces the variability in membrane permeability. When transport across the membrane was essential for expression, this optical treatment increased mean expression level by ~6-fold and reduced expression variability by nearly two orders of magnitude. These results demonstrate membrane engineering may enable essential steps toward cell-like synthetic systems. The experimental platform described here provides a means of understanding controlled transport motifs in individual cells and groups of cells working cooperatively through cell-to-cell molecular signaling.

## Introduction

Cell-free gene expression using purified components or cell extracts has become a viable platform for synthetic biology (Karim and Jewett, 2016; Moore et al., 2018; Pardee et al., 2016; Shin and Noireaux, 2012; Siegal-Gaskins et al., 2014). A broader goal is the realization of more complex cell-free systems (Perez et al., 2016) that may approach cell-like capabilities (Scott et al., 2016). However, these aspirations are stymied by highly variable behavior from identical cell-free expression reactors, especially at cell-relevant reactor volumes (Boreyko et al., 2017; Caveney et al., 2016; Hansen et al., 2015; Norred et al., 2018). Even in simple, single-gene, expression experiments, protein concentrations may vary by more than an order of magnitude across a population of identically constructed reaction chambers (Nourian and Danelon, 2013; Saito et al., 2009). For cell-free expression confined in lipid vesicles, a portion of this variability may emerge from vesicle-to-vesicle variation in the permeability of the membrane (Nishimura et al., 2014a) to molecular species (ions, amino acids, etc.) that may lead to variability in protein production. A small fraction (<10%) of lipid vesicles produce much more protein than the average vesicle because they are naturally permeable to molecular species that support expression (Nishimura et al., 2014a) (Figure 1A). As a result, producing vesicles with more uniform transport properties is an essential step in achieving more uniform expression behavior in synthetic gene expression platforms (Figure 1B).

**Figure 1.**
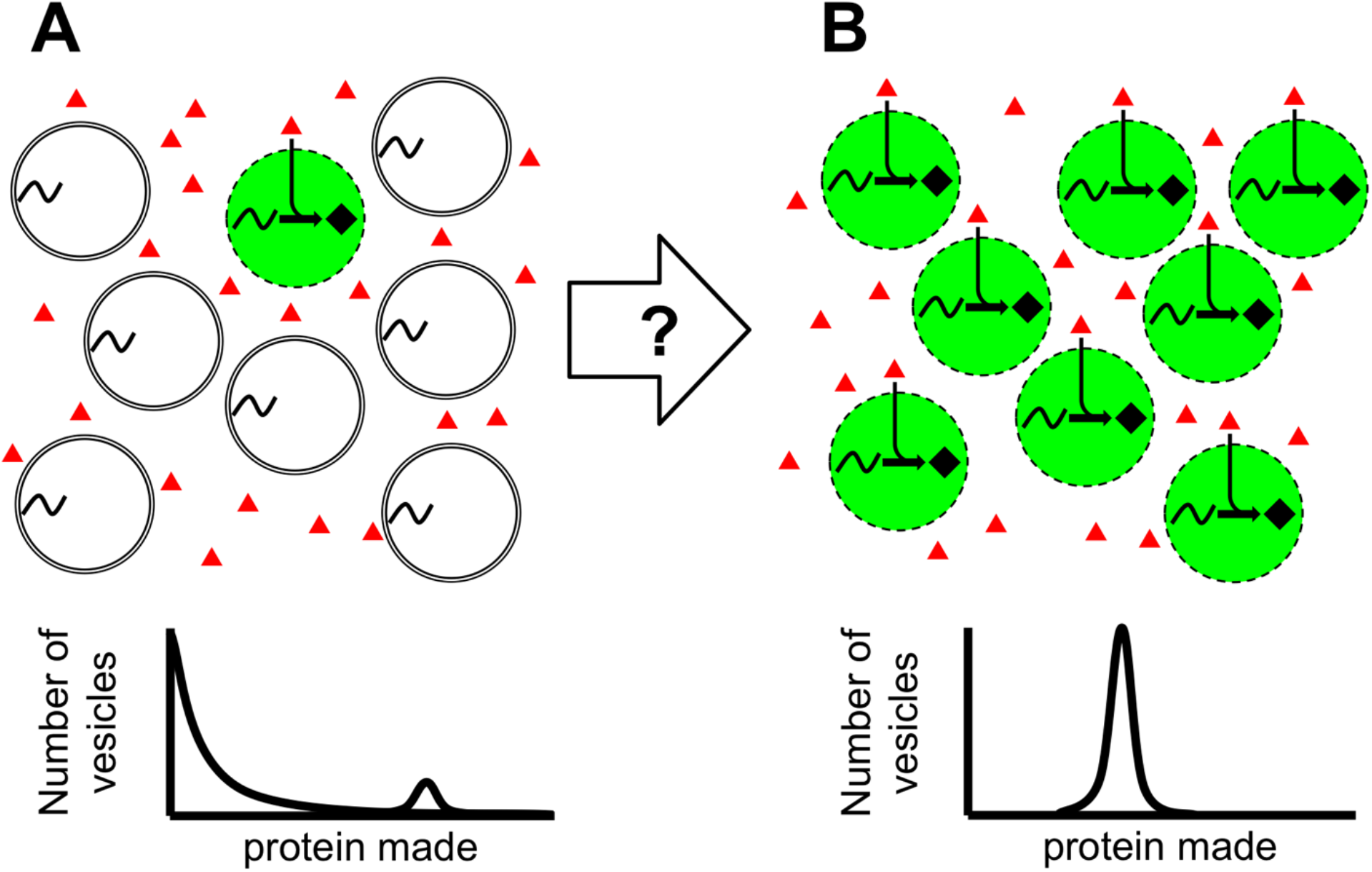
Engineering membranes for uniform protein production. (A) (Top) A population of vesicles where only a few vesicles are permeable to resources essential for expression (red triangles) in the outer solution and thus able to make protein (black diamonds and green background). (Bottom) The result is a highly skewed protein population distribution where most vesicles make no protein and a few make large amounts of protein. (B) A process to make more vesicles permeable to protein expression resources (Top) would result in a more uniform protein population distribution (Bottom).

Recent work suggests that optical exposure induces membrane pore formation (Itri et al., 2014) that may lead to uniform permeability of lipid vesicles (Figure 1B). Such photosensitizing processes have been employed to permeabilize membranes to trigger drug release (Massiot et al., 2019), or to inactivate membrane bound proteins (Rokitskaya et al., 2015). Studies show that optical exposure produces oxidative species (Mertins et al., 2014) that result in chemically modified lipid-tails and increase membrane permeability (Bacellar et al., 2018).

Here we show that for cell-free expression in lipid vesicles, optically-induced permeabilization leads to relatively uniform transport of gene expression resources across the membrane, yet maintains encapsulation of the core expression machinery (e.g. ribosomes and DNA) within the vesicles. In experiments where membrane transport was required for expression activity, this optical treatment led to ~6-fold greater mean protein expression with nearly two orders of magnitude less vesicle-to-vesicle variability in expression level compared to experiments lacking the treatment. For cell-free expression in vesicles, these findings have important implications for the comparison of flow cytometry (minimal optical exposure) to microscopy (more optical exposure) experiments. Importantly, these results demonstrate membrane engineering may enable essential steps toward cell-like synthetic systems with controlled transport of molecules across boundaries for energy exchange, sensing, and communication. As a result, such cell-free experimental platforms provide a viable path for understanding these controlled transport motifs in individual cells and groups of cells working cooperatively through cell-to-cell molecular signaling.

## Results and Discussion

To study how membrane permeability affected gene expression, we tracked cell-free expression of Yellow Fluorescent Protein (YFP) confined in POPC vesicles (Figure 2A; Methods). In these experiments, expression resources encapsulated within the vesicles were deliberately diluted to ensure that transport across the membrane was essential to expression activity (Methods). Each vesicle contained a population of fluorescent molecules (AF647 conjugated to transferrin) captured at the time of vesicle formation to serve as a volume marker that aided in vesicle identification (Figure 2B, top). To ensure that even vesicles that expressed little or no YFP were found, for all experiments the t=2 hour image of the AF647 was used to locate 100-200 regions of interest (ROIs) that indicated the location of individual vesicles possessing distinct boundaries with minimal overlap with neighboring vesicles. The t=2 hour measured YFP fluorescence from these ROIs (Figure 2B, bottom) provided a random sample of the amount of protein expressed in individual vesicles across the population (Figure 2B, bottom inset). Imaging was performed using a Zeiss, LSM 710 confocal laser scanning microscope. Z-stacks of 20 slices were taken during every imaging event for 2 hours. Each slice was 512 x 512 pixels (pixels measured 0.81 μm x 0.81 μm). Vesicles were illuminated with 405 nm, 488 nm, and 633 nm lasers. Laser powers were 6.5 mW, 6.1 mW, and 1.67 mW, respectively. During each image capture, each pixel was illuminated by all three lasers for 37.9 μs.

**Figure 2.**
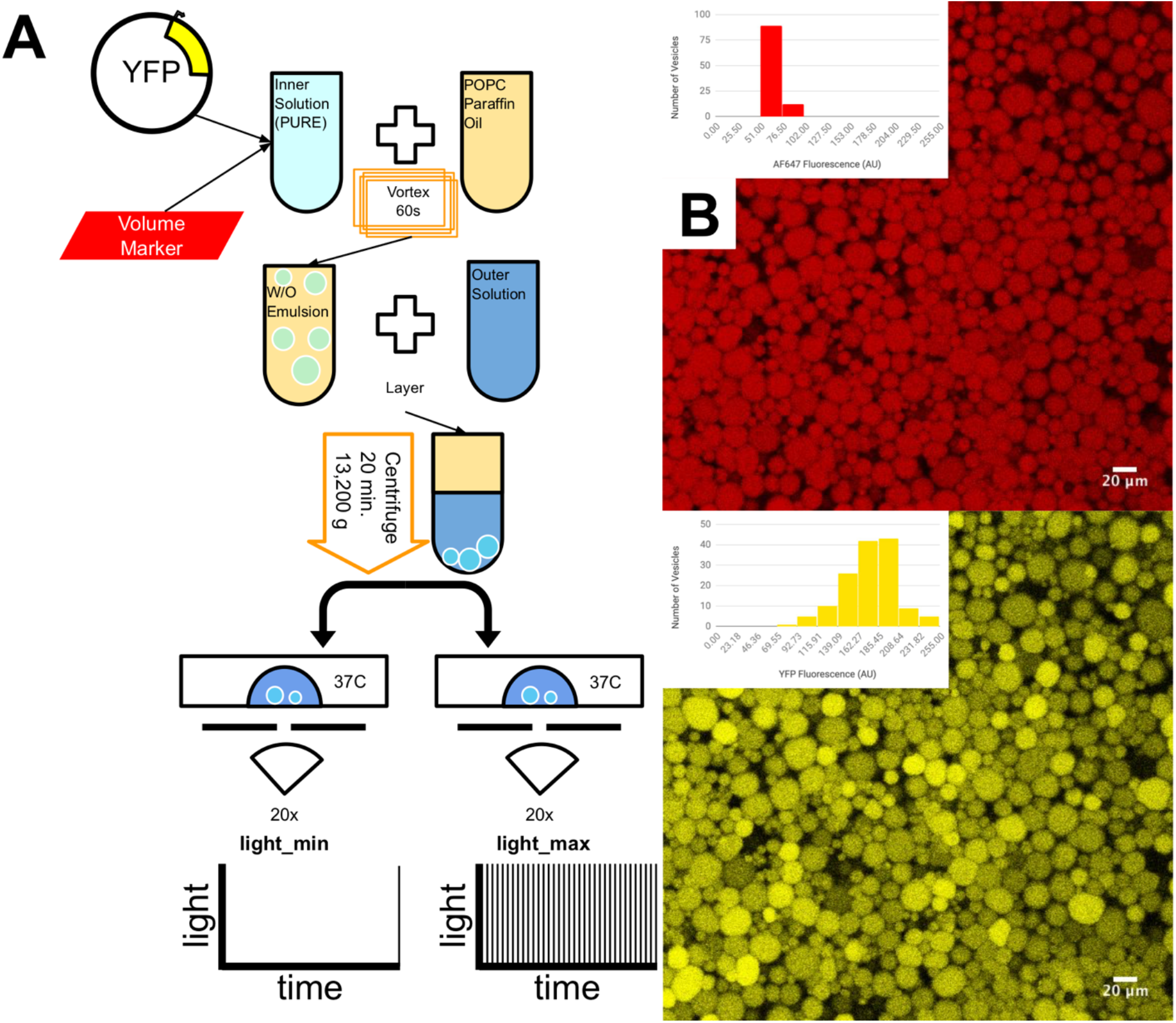
Vesicle production and imaging protocol. (A) Vesicle production via the emulsion-transfer method. YFP expressing plasmid and a fluorescent volume marker (AF647) were encapsulated in lipid vesicles. The vesicles were imaged on a confocal microscope using either a minimal light exposure (bottom left) or maximal light exposure (bottom right) protocol. (B) Images of YFP and AF647 fluorescence after 2 hours of gene expression for a control experiment where membrane transport was not required for robust expression. Insets show the distributions of fluorescent intensities.

We performed experiments using two different imaging protocols. In the first protocol (light_min; Figure 2A, bottom left), we imaged the vesicles only once at t=2 hours after expression activity ceased (Caveney et al., 2016; Karig et al., 2013; Sun et al., 2013). In this protocol, the vesicles were exposed to a minimal amount of light, much as they would be in a flow cytometry experiment. In the second protocol (light_max; Figure 2A, bottom right), we imaged each vesicle once every 3 minutes for the entire 2-hour duration of the experiment. Note that in the light_min protocol there was minimal photobleaching, but that both YFP and AF647 experienced significant photobleaching (18.0% and 32.5%, respectively; SI Figure 1) due to the constant illumination in the light_max protocol.

The AF647 images from both protocols show numerous intact vesicles with distinct boarders (Figure 3A and B). Aside from the photobleaching inherent to light_max (Figure 3C), the distribution of AF647 across the population of vesicles was similar for both protocols (SI Figure 2). In contrast, the expression behavior, as indicated by the YFP images (Figure 3D and E), was vastly different for the two protocols. The light-min protocol resulted in a skewed population (adjusted Fisher–Pearson standardized moment coefficient (g1) =6.25) where most vesicles (76.6%) had little or no YFP expression, 20.3% made some detectable amount of YFP, and a small number (3.1%) made much more YFP than the average. These results are consistent with previous reports of POPC vesicles made by the emulsion-transfer method expressing protein with the PURE system (Nishimura et al., 2014a) that show ~10% of vesicles are naturally permeable to small molecules necessary for gene expression. Conversely, the majority of vesicles are impermeable and may produce little or no protein if they are lacking essential expression resources encapsulated within the vesicle (Nishimura et al., 2014a).

**Figure 3.**
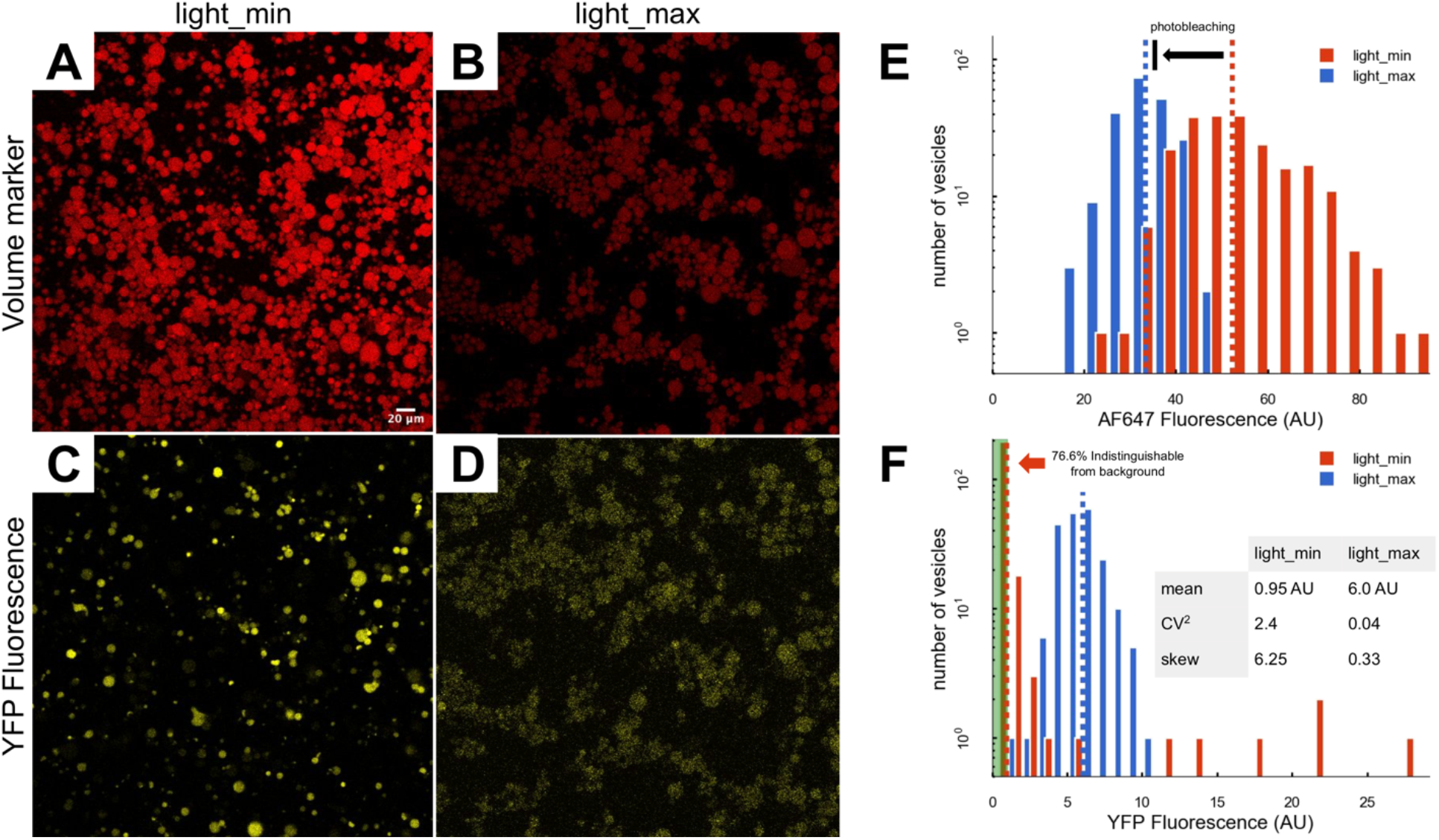
Effects of the light_min and light_max protocols on gene expression behavior. The t=2 hour image of AF647 using (A) light_min and (B) light_max protocols. (C) The t=2 hour distributions of AF647 fluorescence for light_min (red) and light_max (blue) protocols. The dashed vertical lines indicate the means of the two distributions. The arrow shows the expected shift in the mean due to photobleaching in the light_max protocol. The t=2 hour image of YFP using the (D) light_min and (E) light_max protocols. (F) The t=2 hour distributions of YFP for light_min (red) and light_max (blue) protocols. The dashed vertical lines indicate the means of the two distributions.

In contrast to the light_min protocol, the light_max protocol resulted in all vesicles expressing measurable levels of YFP with a much less skewed distribution (g1=0.33). As a result, even without correcting for photobleaching (Figure 3F; SI Figure 1), the mean YFP expression level was ~6-fold greater for light_max compared to light_min. Further, the light_max protocol decreased expression variability (measured using CV^2^ = YFP variance/YFP mean) by nearly two orders of magnitude (CV^2^=2.4 (light_min); CV^2^=0.04 (light_max)).

The clear implication of the experimental results is that the light_max protocol permeabilized the vesicle membranes. Although not previously reported in gene expression studies, such photosensitive permeabilization of lipid membranes has been reported in other contexts (Figure 4A). These reports (Itri et al., 2014; Mertins et al., 2014) indicate that light exposure creates short-lived, excited molecules that react with molecular oxygen to produce singlet oxygen.

**Figure 4.**
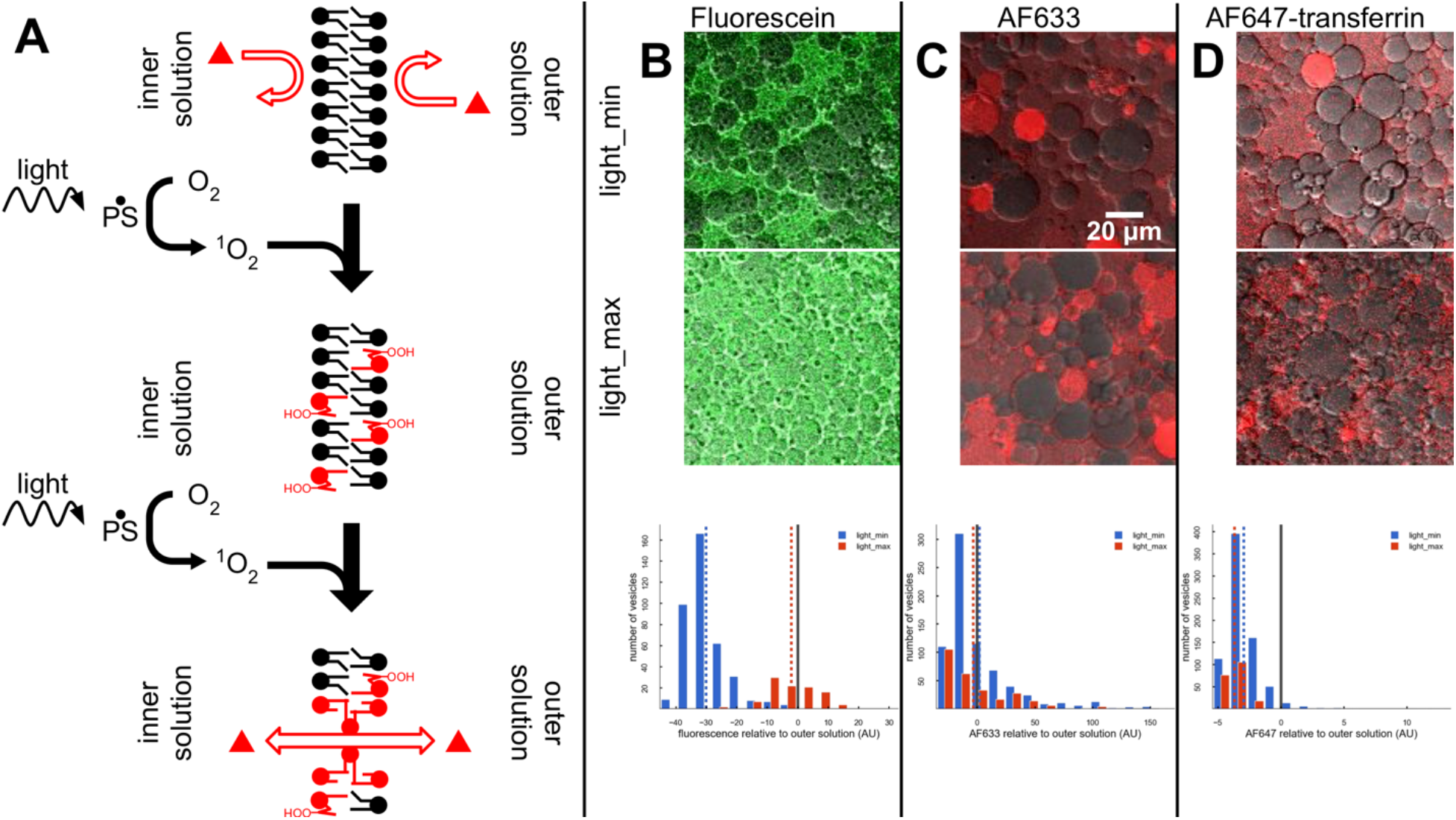
Permeabilization of POPC membranes. (A) Light exposure excites photosensitive species which produce singlet oxygen that reacts with the unsaturated tails of POPC to form alcohol and ketone groups, leading to permeabilization of the membrane (De Rosa et al., 2018). (B-D) Transport of fluorophores from outer solution into the vesicles after 2 hours of light_min (top) or light_max (middle) imaging protocol for (B) Fluorescein, (C) AF633, and (D) AF647. The light_max protocol enables Fluorescein transport into the vesicles (B, bottom), but has little effect on AF633 (C, bottom) or AF647 (D, bottom) transport into vesicles. In (B-D, bottom) the dashed vertical lines indicate the means of the populations. The solid black vertical lines indicate concentration of fluorophore in the outer solution.

Singlet oxygen reacts with double bonds found on lipid tails, resulting in a shift of the double bond by one carbon, and most importantly, generates lipid hydroperoxide species. The double bond shift increases the area per lipid molecule (Mertins et al., 2014), decreases membrane thickness (Itri et al., 2014) and the elastic moduli (Weber et al., 2014), but does not increase membrane permeability (Weber et al., 2014). However, the hydroperoxides produce alcohols and ketones on lipid tails, as well as truncated lipid tails capped with an aldehyde that form stable pores in the membrane (De Rosa et al., 2018) (Figure 4A, bottom).

While the gene expression data suggest optically-induced pore formation in the vesicle membranes, they also imply molecular selectivity in transport through the pores. Most importantly, significant levels of YFP were only seen in vesicles. Furthermore, the drop in AF647-transferrin fluorescence in the light_max protocol was consistent with photobleaching and showed no evidence of significant loss due to leakage from the vesicles. Accordingly, it seems that the larger expression resource molecules (plasmid DNA, ribosomes, RNAPs) and the proteins (transferrin, YFP) remained encapsulated, while smaller molecular species (small ions, nucleotides, amino acids) were able to cross the membrane.

To characterize the size-electivity of the membrane pores, we imaged populations of vesicles with various fluorophores in the outer solution (Figure 4B-D). The fluorophores were chosen to have molecular weights larger than small ions, but in the range of amino acids (~110 Da), nucleotides (~650 Da), and proteins (>20 kDa). Fluorescein (~332 Da), AF633 (~1.2 kDa), and AF647 conjugated to transferrin (~80 kDa) were added to the outer solution and the vesicles were imaged with both the light_min (Figure 4B-D, top) and light_max (Figure 4B-D, middle) protocols. The light_min protocol resulted in little or no diffusion of Fluorescein or AF647 into vesicles (Figure 4B and D, bottom). In contrast, light_max imaging enabled diffusion of Fluorescein across the membrane, equalizing the vesicle and outer solution concentrations (Figure 4B, bottom). Light_max imaging resulted in little change in the populations of AF633 and AF647-transferrin within the vesicles. However, even with very little optical exposure, AF633 was found in many vesicles, often at concentrations greater than the outer solution (Figure 4C, bottom). This initial loading of fluorophore into vesicles was seen from the very first image in the light_max protocol and was unrelated to optical exposure. It seems likely that AF633 may become encapsulated in some vesicles during the vesicle synthesis process. These fluorophore transport and the gene expression results indicate that the light_max protocol enabled the transport of small ions, nucleotides, and amino acids, yet kept RNAP (~99 kDa), proteins, ribosomes (~2.7 MDa), and plasmid DNA (~1.58 MDa) encapsulated within the vesicles.

Synthetic biology approaches to controlling gene expression level and variability have focused on genetic circuits. However, the central feature of cell-free synthetic biology is the ability to define the environment by manipulating confinement volume (Caveney et al., 2016), degree of macromolecular crowding (Norred et al., 2018), and the composition of cell extract (Garcia et al., 2018). Likewise, the results reported here show that membrane engineering is a viable approach to control expression behavior and demonstrate a simple optical treatment that greatly diminishes the variability in the permeability of POPC vesicle membranes. Looking ahead, such controlled transport across membranes is essential for energy exchange, sensing, and communication that lie at the heart of many complex cellular functions. The intriguing implication of this study is that membrane engineering may enable cell-like synthetic systems with similar levels of functionality. As a result, such cell-free experimental platforms provide a viable path both for the realization of cell-free synthetic biology applications and for understanding these controlled transport motifs in individual cells and groups of cells working cooperatively through cell-to-cell molecular signaling.

## Methods

### Vesicle Preparation

Vesicle were made using the oil-in-water emulsion-transfer method (Noireaux and Libchaber, 2004; Pautot et al., 2003) (Figure 2A). This method encapsulated a protein expressing inner solution in vesicles separated from an osmotically balanced outer solution. The inner solution was prepared using 10 μL Solution A and 7.5 μL Solution B of the PURExpress In Vitro Protein Synthesis Kit from New England Biolabs; 5 μL of sucrose solution (1 M); 0.25 μL of Transferrin-AlexaFluor 647; 0.125 μL of RNAsin (40 U/μL); 0.418 μL (1.67 nM) of YFP encoding pEToppYB plasmid (Nishimura et al., 2014b) (200 ng; 478.2 ng/μL stock); and nuclease-free water to bring the total volume of solution to 30 μL. The inner solution was vortexed in 330 μL of paraffin oil containing 30 mg of 1-palmitoyl-2-oleoyl-glycero-3-phosphocholine (POPC) for 60 seconds. The resulting emulsion was layered above the outer solution and centrifuged at 13,000 g for 20 minutes at room temperature. The low concentration inner reactions were made by diluting Solution A and Solution B with nuclease-free water to 1/3 their standard concentrations.

### Outer Solution Preparation

The outer solution for vesicles was mixed from frozen stocks before each experiment. 1.5 μL Amino acid solution, 11.3 μL of ATP (100 mM), 7.5 μL of GTP (100 mM), 0.75 μL of CTP (500 mM), 0.75 μL of UTP (500 mM), 1.8 μL of spermidine (250 mM), 3.75 μL of creatine phosphate (1 M), 4.5 μL of Dithiothreitol (100 mM), 0.75 μL of Folinic Acid (0.5 M), 24 μL of potassium glutamate (3.5 M), 11.3 μL of magnesium acetate (0.5 M), 30 μL of HEPES (1 M), 60 μL of glucose (1 M), and 141.8 μL of autoclaved type I pure water for a total volume of 300 μL.

### Vesicle Imaging

The pellet of vesicles was collected with 100 μL of the outer solution and pipetted onto a no. 1.5 glass bottom petri dish. The lid was placed on the petri dish to minimize airflow and evaporation of the 100 μL outer solution and vesicle drop. Two different protocols were followed for imaging: light_max and light_min. For the light_max protocol the petri dish was placed on a Zeiss LSM710 confocal scanning microscope with an incubation chamber warmed to 37°C and imaged every 3 minutes in a z-stack with a 20x air objective. Vesicles were imaged with three lasers: a 405 nm, 6.5 mW laser; YFP was excited with a 488 nm, 6.1 mW laser and fluorescent emission was collected from 515-584 nm; and AF647 was excited with a 633 nm, 1.67 mW laser and fluorescent emission was collected from 638-756 nm. Z-stacks were made of 20 slices at 1 μm intervals, and the aperture for each slice was 1.00 Airy Units (open enough to allow ~1.5 μm depth of light). The time the vesicles sat on the microscope before imaging was minimized (less than 15 minutes), allowing for imaging for most of the duration of protein expression. For the light_min protocol, the petri dish with vesicles was placed in a dark incubator at 37°C for 2 hours. It was then imaged once on the confocal microscope with the same settings as the light_max protocol.

### Data Acquisition and Analysis

Average fluorescent intensity and diameter were measured with the FIJI TrackMate (Tinevez et al., 2017) (v3.8.0) plugin. TrackMate found spots with an estimated blob diameter of 10 μm using the Laplacian of Gaussian detector. Spots that were found with an estimated diameter <5 μm, >19 μm, or contrast <0 were removed from the data set. We used the simple Linear Assignment Problem (LAP) tracker to link spots across z-stacks in time to create traces. Traces that had missing frames, traveled >5μm between frames, or tracked for <45 of the 60 frames were removed from the data set. Fluorescent concentrations and estimated diameters were measured for each vesicle at each time point. The estimated diameter varied slightly between frames, so the diameter used for each vesicle was taken to be the average of the estimated diameters over the entire time trace. Protein concentration (in arbitrary units) was found by multiplying the average fluorescent intensity by the volume.

### Determining Vesicles Indistinguishable from Background

Background ROIs were determined from the AF647 (volume marker) channel. The gene expression intensities of these ROIs were measured in the YFP channel. A sample of vesicles with clearly defined edges in the AF647 channel, but no clear edges in the YFP channel, were measured for their YFP intensity. This value was slightly above the background value measured and was used as a cut-off for vesicles that were indistinguishable from background fluorescence.

## Acknowledgements

This research was conducted at the Center for Nanophase Materials Sciences, which is a DOE Office of Science User Facility. P.M.C., R.N.D, S.E.N., and W.T.M also acknowledge Graduate Fellowships from the Bredesen Center for Interdisciplinary Research and Graduate Education, University of Tennessee, Knoxville. The authors would like to thank Osaka University and Dr. Tetsuya Yomo for providing pEToppYB plasmid, and Dr. Maike Hansen and the Dr. Leor Weinberger laboratory for useful discussions.

## Author Contributions

Conceptualization, P.M.C, S.E.N, C.P.C, and M.L.S; Methodology, P.M.C, S.E.N, C.P.C, and M.L.S; Formal Analysis, P.M.C, M.L.S.; Investigation, P.M.C, R.M.D., and W.T.M; Writing – Original Draft, P.M.C, W.T.M, and M.L.S; Writing – Reviewing & Editing, P.M.C and M.L.S; Visualization, P.M.C; Supervision, C.P.C, M.L.S; Funding Acquisition, M.L.S.

