## Supplementary material for "Controlling Cell-Free Gene Expression Behavior by Tuning Membrane Transport Properties": SI Figure

### Supplemental Figures

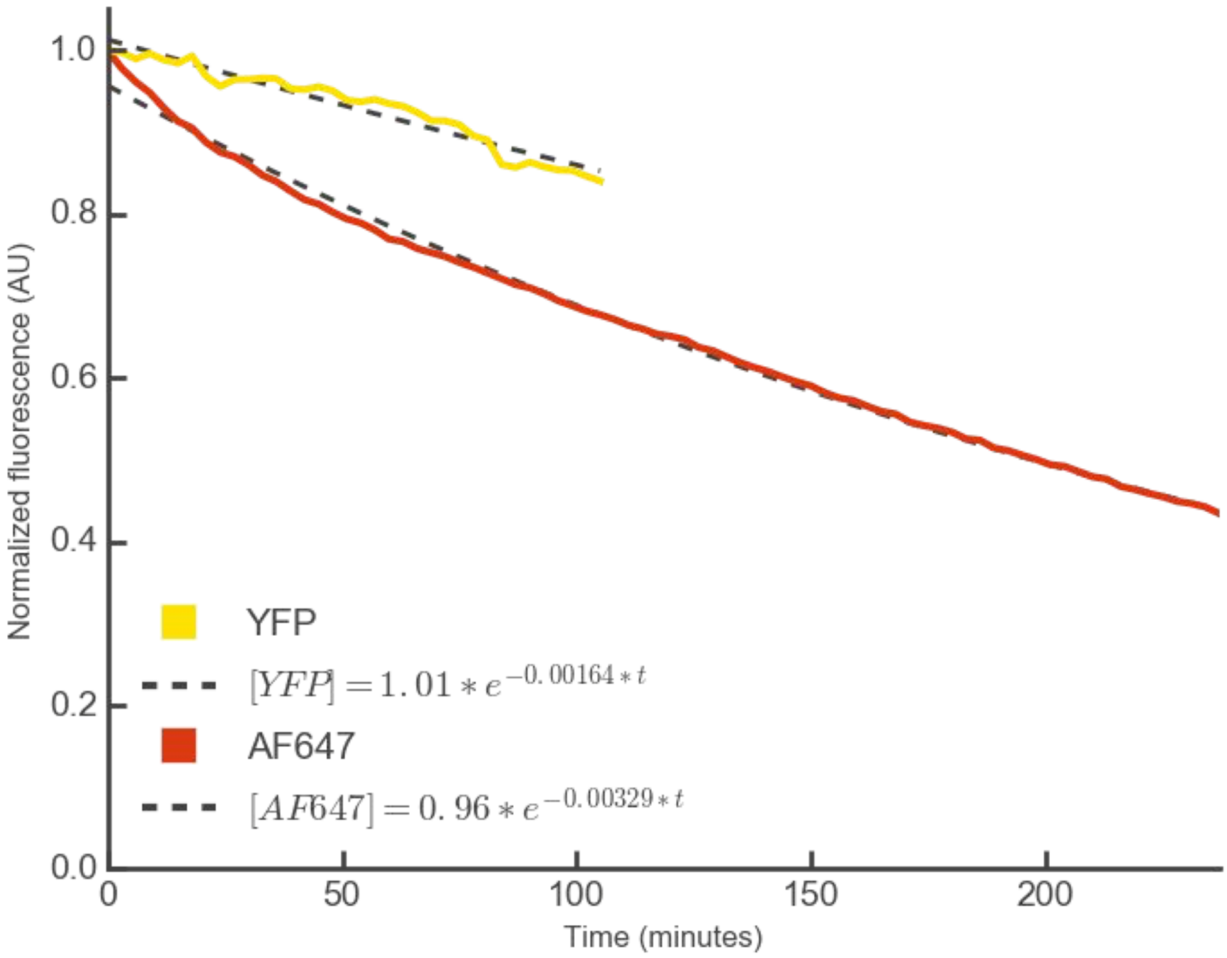


**SI Figure 1: Photobleaching rate of YFP and AF647 during the light_max protocol.** A standard vesicle reaction was prepared and left for two hours in the dark to finish making protein. Vesicles were then imaged using the light_max protocol to measure the YFP and AF647 photobleaching rates. The photobleaching half-life was 420 minutes for YFP and 211 minutes for AF647. After two hours, the light_max protocol reduced YFP fluorescence by 18% and AF647 fluorescence by 32.5%.

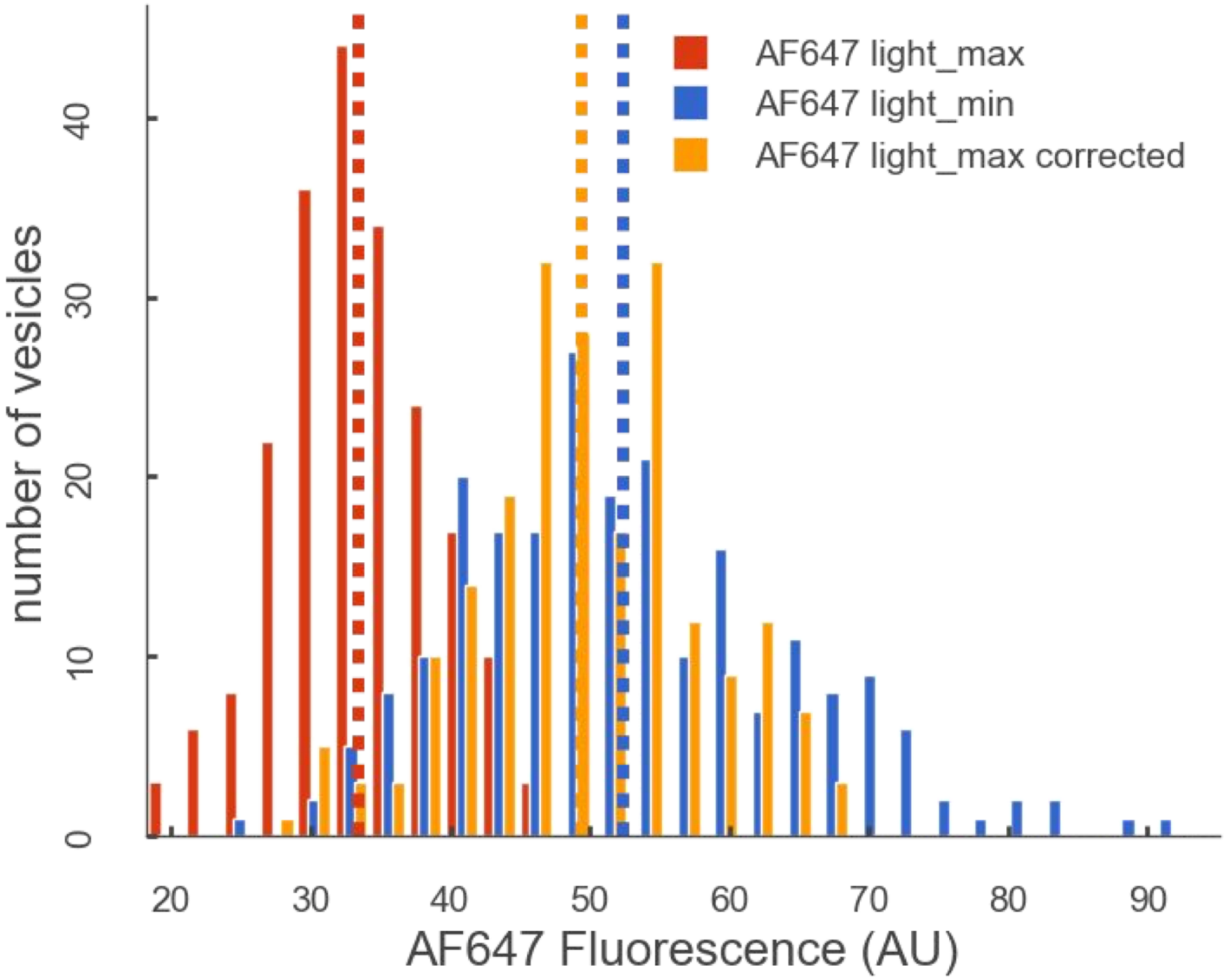


**SI Figure 2: Light_max effect on AF647 fluorescence distribution.** To compare AF647 distributions from light_min (blue) and light_max (red) protocols, the light_max distribution was corrected for photobleaching (orange) by multiplying by a bleaching gain factor (1.48). The dashed vertical lines indicate the means of the populations.
